# COVID-MVP: an interactive visualization for tracking SARS-CoV-2 mutations, variants, and prevalence, enabled by curated functional annotations and portable genomics workflow

**DOI:** 10.1101/2022.06.07.493653

**Authors:** Muhammad Zohaib Anwar, Ivan S Gill, Madeline Iseminger, Anoosha Sehar, Kenyi D Igwacho, Khushi Vora, Gary Van Domselaar, Paul M. K. Gordon, William WL Hsiao

**Affiliations:** Centre for Infectious Disease Genomics and One Health, Faculty of Health Sciences, Simon Fraser University, Burnaby, BC, Canada; University of British Columbia, Vancouver, BC, Canada; National Microbiology Laboratory, Public health Agency of Canada, Winnipeg, MB, Canada; Centre for Health Genomics and Informatics, Cumming School of Medicine, University of Calgary, Calgary, AB, Canada

**Keywords:** SARS-CoV-2, mutations, genomic variants, genomic surveillance, functional annotation, visualization, genomic epidemiology

## Abstract

The SARS-CoV-2 pandemic has reemphasized the importance of genomic epidemiology to track the evolution of the virus, dynamics of epidemics, geographic origins, and the emerging variants. It is vital in understanding the epidemiological spread of the virus on global, national, and local scales. Several analytical (bioinformatics) resources have been developed for molecular surveillance. However, a resource that combines genetic mutations and functional annotations on the impact of these mutations has been lacking in SARS-CoV-2 genomics surveillance. COVID-MVP provides an interactive visualization application that summarizes the mutations and their prevalence in SARS-CoV-2 viral lineages and provides functional annotations from the literature curated in an ongoing effort, Pokay. COVID-MVP is a tool that can be used for routine surveillance including spatio-temporal analyses. We have powered the visualization through a scalable and reproducible genomic analysis workflow nf-ncov-voc wrapped in Nextflow. COVID-MVP allows users to interactively explore data and download summarized surveillance reports. COVID-MVP, Pokay, and nf-ncov-voc are open-source tools available under the Massachusetts Institute of Technology (MIT) and GPL-3.0 licenses. COVID-MVP source code is available at https://github.com/cidgoh/COVID-MVP and an instance is hosted at https://covidmvp.cidgoh.ca.

## 1. Introduction

The first case of COVID-19 was officially confirmed in December 2019 and some studies even suggest earlier evidence [1]. In the following three months, hundreds of other countries reported at least one confirmed case of Covid-19, and the World Health Organization (WHO) formally declared it a global pandemic on March 11th, 2020 [2]. Since then, cases started to increase exponentially, and currently (June 2022), more than 500 million cases have been reported, with more than 6 million deaths worldwide [3]. The first genome sequence of severe acute respiratory syndrome coronavirus 2 (SARS-CoV-2)—the virus which causes COVID-19—was released on January 10th, 2020 [4]. Facilitated by Whole Genome Sequencing (WGS) and data analytics (bioinformatics) methodologies, an unprecedented amount of viral genomic data has been generated and analyzed during this pandemic--more than 10 million viral genome sequences generated worldwide have been deposited to the GISAID international SARS-CoV-2 sequence repository. These efforts have demonstrated the feasibility and benefits of near-real-time genomic surveillance and genomic epidemiology of pathogens. Genomic epidemiology has substantial advantages over conventional methods as sequence data produced can be used for more accurate detection and characterization of genomic variants [5,6]. The first SARS-CoV-2 variant identified to substantially increase disease transmissibility was first detected in the later months of 2020 [7]; since then, the WHO has classified variants as Variants of Concern (VOC), Variants of Interest (VUI), and Variants under Monitoring (VUM) based on their level of assessed risk [8]. Numerous publicly available genomic surveillance platforms have been developed to understand the evolution and transmission of SARS-CoV-2; for example, Nextstrain [9] uses time-scaled phylogenies to connect source and transmission chains of Covid-19 outbreaks [10,11]. CoVizu [12] reconstructs robust evolutionary and epidemiological relationships among SARS-CoV-2 genomes. It provides rapid analysis and visualization of the global diversity of SARS-CoV-2.

Understanding the functional significance of the mutations harbored by a variant is important to assess its impact on transmissibility, disease severity, immune escape, and effectiveness of vaccines and therapeutics. [13]. Although, it was initially estimated that the mutation rate of the SARS-CoV-2 genome was approximately one mutation per week, with a Non-synonymous/Synonymous mutation ratio (Ka/Ks) of 1.008 [14,15], a more recent study [16] suggested the mutation rate is at least 49–67% higher based on the rate of appearance of variants in sampled genomes. These mutations may or may not change the biological structure (and functional impact) of a virus which is why a near-real-time visualization of emerging mutations is also a vital aspect of genomic epidemiology. A few visualization dashboards and analytical workflows have been developed to track the prevalence of mutations of concern such as the COVID-19 virus mutation tracker system (CovMT) [17] which shows the evolution of the mutational landscape of SARS-CoV-2. CovMT summarises mutations with respect to geographical location, date of sampling, and disease severity (when this information is available). Similarly, CovRadar [18] is specifically designed for molecular surveillance of the spike gene of SARS-CoV-2. It focuses on mutations and provides easy access to frequencies and spatio-temporal distributions from global sample collections. Outbreak.info is another web-based visualization dashboard that provides a similar heat map-based user interface for investigating the mutational content of SARS-CoV-2 lineages [19]. However, an application that integrates lineage information with genetic mutations and functional annotations on the impact of these mutations has been lacking in SARS-CoV-2 genomics surveillance. Here, we present a visualization-focused application, COVID-MVP, powered by a manually curated functional annotation *pokay and* the high-throughput genomic analysis workflow *nf-ncov-voc*. COVID-MVP is an open-source project to provide an interactive tool to visualize the mutational content and functional significance of SARS-CoV-2 genomes.

## 2. Results

We have developed an interactive framework for visualizing and reporting mutations in SARS-CoV-2 variants. This framework is composed of three stand-alone yet connected components; an interactive visualization (COVID-MVP), a manually curated functional annotation database (*pokay*), and a genomic analysis workflow (*nf-ncov-voc)*. COVID-MVP provides (i) an interactive heatmap to visualize and compare mutations in SARS-CoV-2 lineages (∼230 as of May 15, 2022) classified across different VOCs, VOIs, and VUMs; (ii) mutation profiles including the type, impact, and contextual information; (iii) annotation of biological impacts for mutations where functional data is available in the literature; (iv) summarized information for each variant and/or lineage in the form of a surveillance report; and (v) the ability to upload raw genomic sequence(s) for rapid processing and annotating for real-time classification (see Supp. Table 1 for summarized comparison with similar tools/dashboards). COVID-MVP is developed using the Plotly and Dash frameworks of Python. Pokay is a tool developed to report salient DNA (qPCR primer/probe) or amino acid (epitope) mismatches info against a set of pathogen isolate genomes specifically SARS-CoV-2. Pokay also includes a manually curated functional annotation dataset captured from the available literature. This dataset is an effort to curate all the functions associated with SARS-CoV-2 mutations. nf-ncov-voc is developed in the nextflow [20] framework including python scripts. It can also be integrated into other surveillance tools. This workflow rapidly assigns lineages to SARS-CoV-2 sequences using a manually curated Pango nomenclature system that classifies the global diversity of SARS-CoV-2 into a hierarchy of lineages [21] and identifies mutations. The mutations are then annotated using the functional annotation curated in *pokay* and integrated into a Genome Variant File (GVF) for interactive visualization in COVDI-MVP. Additionally, nf-ncov-voc generates surveillance reports available to download through COVID-MVP. An instance of this tool is presently hosted at https://covidmvp.cidgoh.ca (last accessed May 15th, 2022). Source code for all three components is available as open-source projects (see *Availability of Data and Materials*) under the Massachusetts Institute of Technology (MIT) and GPL-3.0 licenses.

### 2.1 Front-end visualization

The COVID-MVP frontend is implemented almost entirely in Python, using the graphing library Plotly and the visualization framework Dash. The backend *nf-ncov-voc* pipeline generates the mutation data integrated with the functional annotations (see Back-end data analysis section). Two implementations of the front-end COVID-MVP web interface are available: one with the “upload button”, and one that does not support uploading data. Both versions are available for local and cloud deployment and can be cloned from the GitHub repository (see *Availability of Data and Materials*).

#### 2.1.1 Heatmap

As shown in Figure 1, the front-end of COVID-MVP is an interactive web application centered around a heatmap that encodes the frequency of mutations across SARS-CoV-2 lineages. The heatmap has multiple axes, encoding the following attributes: (i) Top: nucleotide position of mutation w.r.t to the first SARS-CoV-2 lineage sampled in Wuhan (GenBank accession MN908947.3; [4]); (ii) Bottom: amino acid position of the mutation within its gene, or if the mutation is in an intergenic region, the number of amino acid positions downstream of the closest gene; (iii) Left: SARS-CoV-2 lineages, with VOC and VOI in bold and italics respectively and (iv) Right: Number of sequences used to generate data per each SARS-CoV-2 lineage.

**Figure 1.**
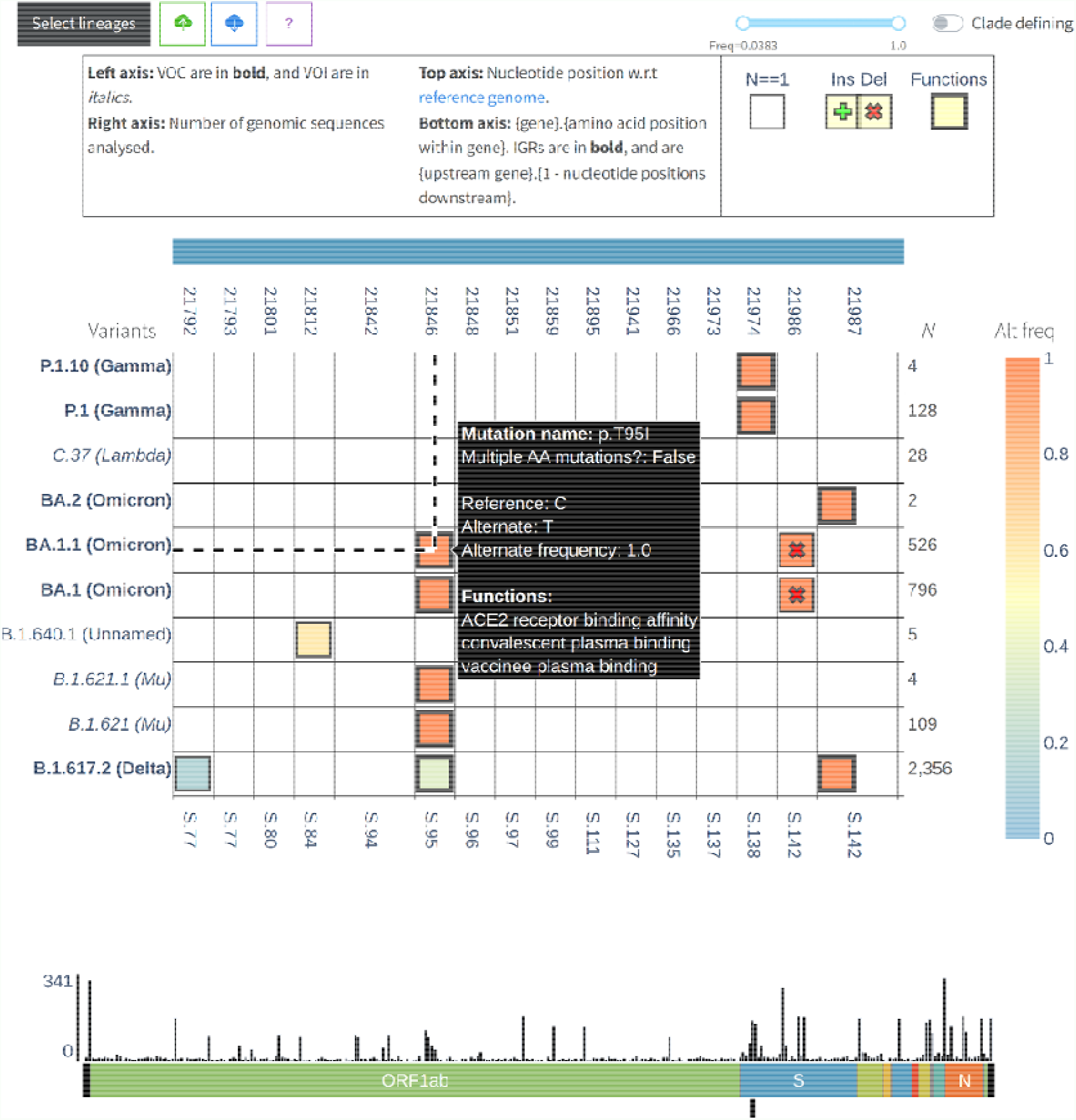
Screenshot of COVID-MVP frontend web interface. The central heatmap encodes mutation frequency across SARS-CoV-2 lineages, with functional information available on hover.

Heatmap cells encoding insertion and deletion mutations are annotated with unique markers, and cells representing mutations with known functions are represented with a thicker border. Users can hover over the heatmap cells for more information on each mutation, and if the mutations have associated functions, users can also click on the cells to open a modal with a description and source of each function, as shown in Figure 2.

**Figure 2.**
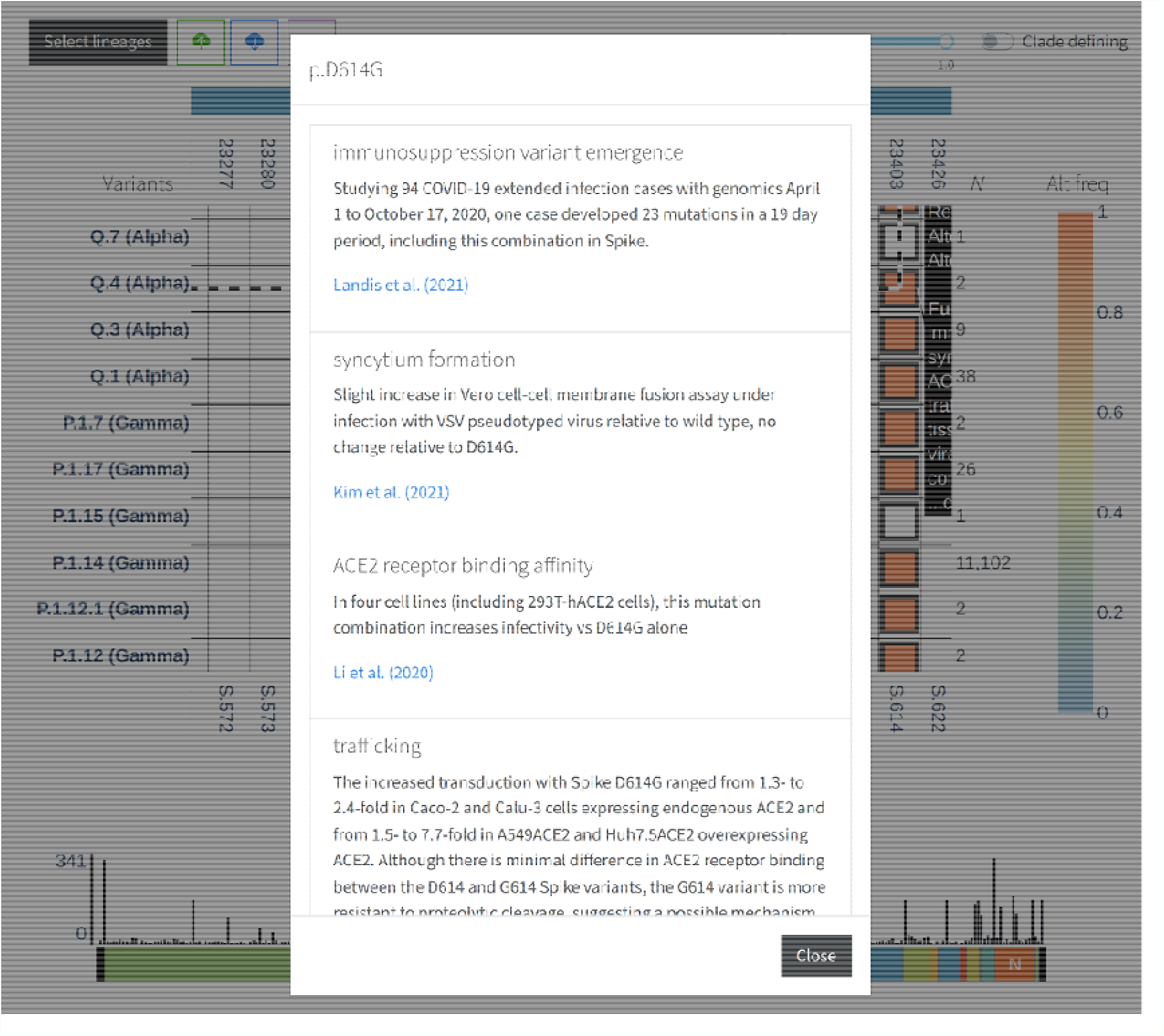
The modal opens when a user clicks on a heatmap cell with functional annotations in COVID-MVP. A description of each function is provided, along with its primary source as a clickable link.

#### 2.1.2 Histogram

Under the heatmap, a histogram encodes the total number of mutations across all visualized lineages per every 100 nucleotide positions w.r.t the reference. The goal of this secondary visualization is to summarize the distribution of mutations across the genome.

#### 2.1.3 Editing the visualization

The frontend visualization can be edited in several ways. A “select lineages” button at the top-left of the application opens a modal that allows users to filter or change the order of lineages visualized in the heatmap. A “clade-defining” switch will enable users to quickly filter out mutations with <25% frequency, based on the “clade-defining” description at Outbreak.info [19]. However, a frequency slider to the left of this switch provides greater granularity for filtering mutations by frequency if necessary.

#### 2.1.4 Downloading/uploading data

An “upload” button to the right of the “select lineages” button allows users to upload a SARS-CoV-2 VCF file, which is processed by the backend *nf-ncov-voc* pipeline, and then visualized in the heatmap alongside the existing data. Following the “upload” button a “download” button is also available that allows users to download a ZIP file containing the surveillance reports of all lineages visualized in the heatmap, as generated by the *nf-ncov-voc* pipeline.

### 2.2 Back-end data analysis

The COVID-MVP backend (*nf-ncov-voc*) is implemented in the Nextflow workflow language with support from Python scripts. Raw sequence data and metadata are downloaded from the Canadian VirusSeq Data Portal for every data release (https://virusseq-dataportal.ca last accessed May 15th, 2022) and processed to visualize at a public instance hosted at https://covidmvp.cidgoh.ca/. The backend workflow consists of four modular sub-workflows that combine in a “plug and play” manner. A detailed data flow diagram is presented in Figure 3.

**Figure 3.**
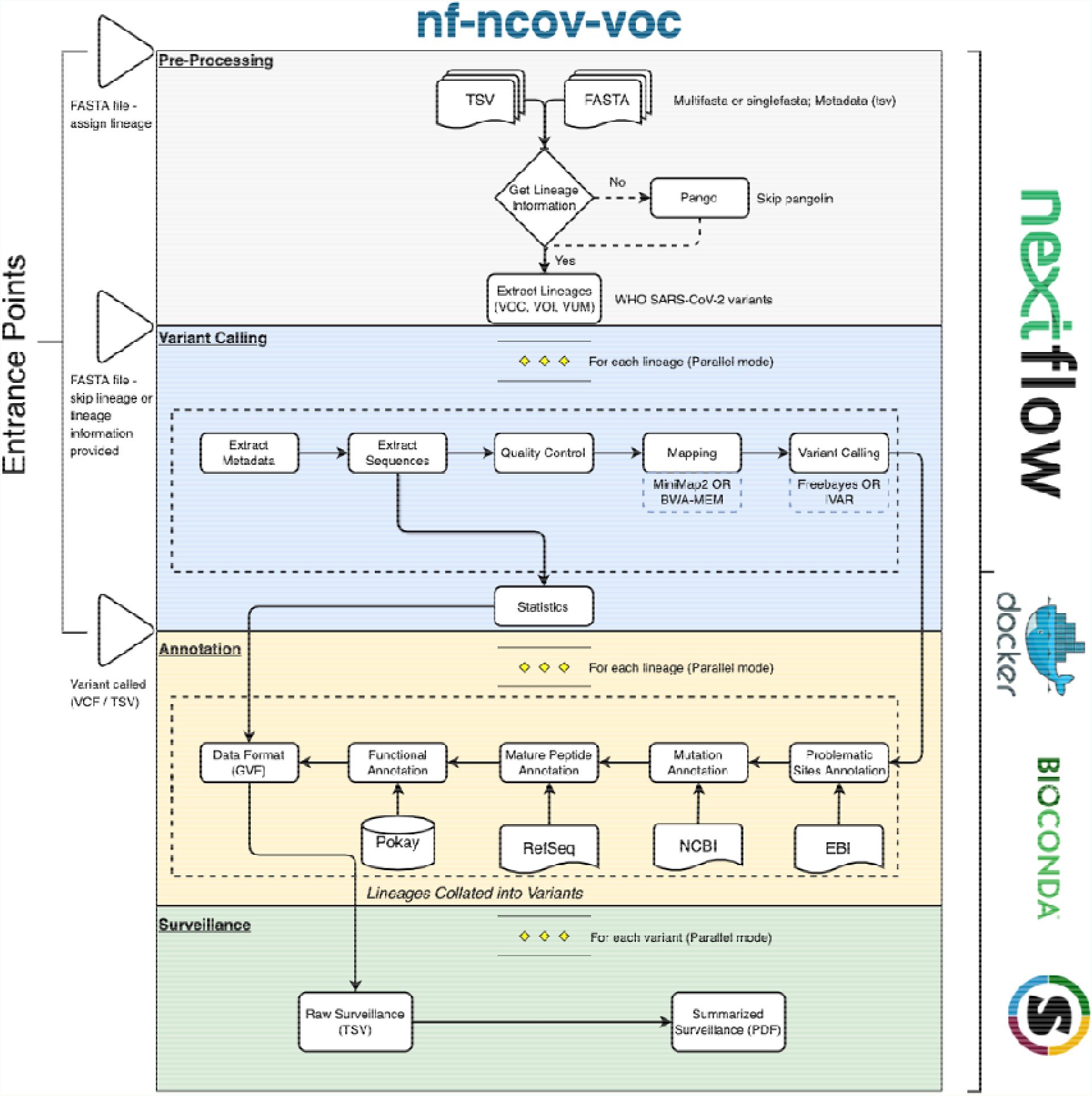
Data flow diagram of genomics workflow

**Figure 4.**
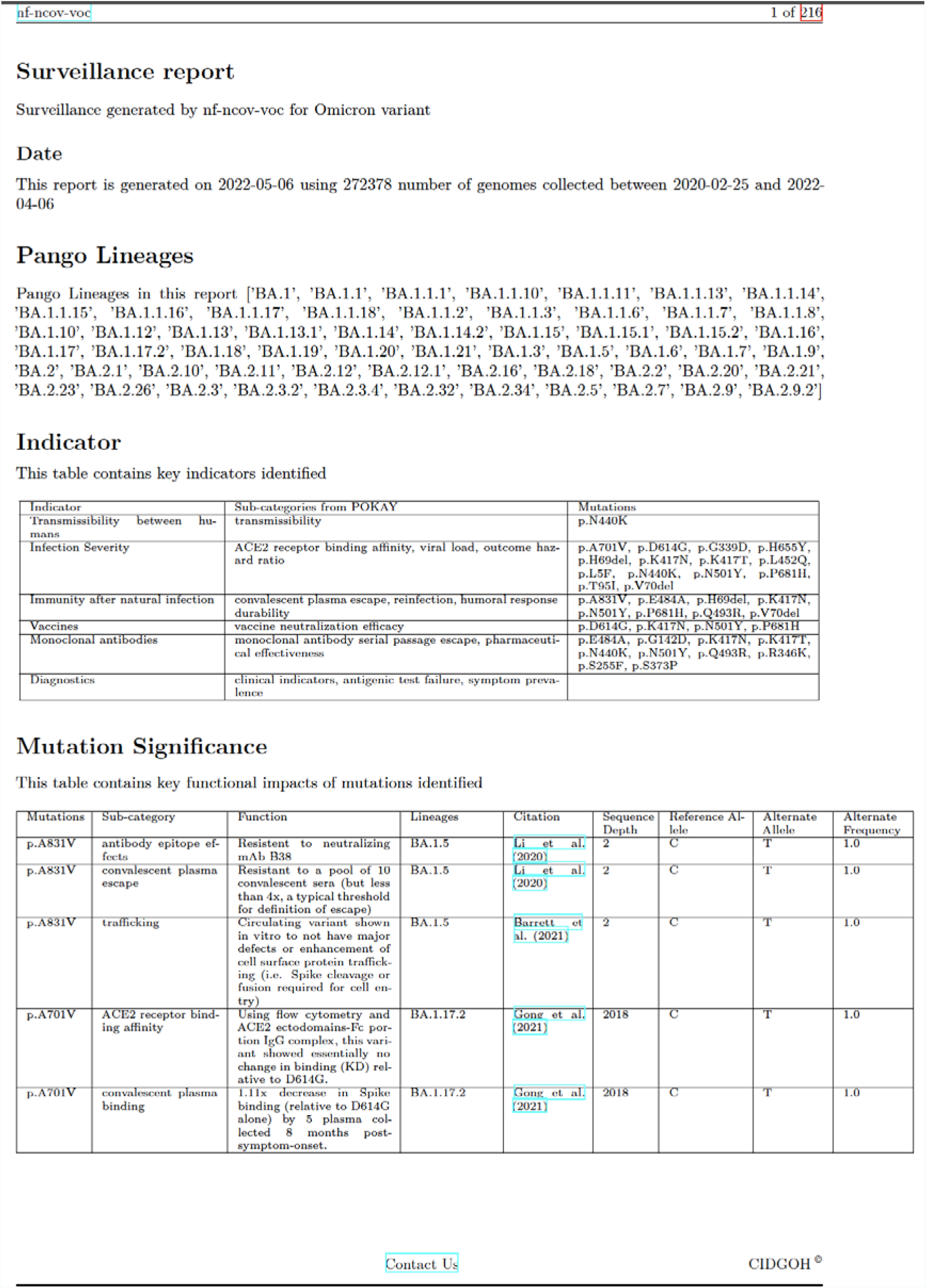
The surveillance report provides insights into the data used in COVID-MVP. It provides information on the number of genome sequences, identified viral (pango) lineages, a table highlighting mutations associated with high-level surveillance indicators, and a table to summarize the significance of each mutation

#### 2.1.1 Pre-processing

SARS-CoV-2 genome sequences in (Fasta format) and accompanying minimal metadata or contextual data (TSV format) are streamed into the workflow. If the lineage information is not available in the metadata file, the sequence file can be used to run Pangolin [21]. Pangolin generates a report file which is then merged with the metadata file using a custom Python script. Alternatively, if the lineage information is already available, sequences and metadata are filtered based on the lineages that belong to VOCs, VOIs, and VUMs. Additional metadata (TSV format) file is curated and updated regularly based on the WHO SARS-CoV-2 variants tracking report https://www.who.int/en/activities/tracking-SARS-CoV-2-variants/ (last accessed May 15th, 2022) to determine the lineages classified as VOC, VOI or/and VUM.

#### 2.2.2 Variant calling

Based on the metadata records, a filtering process is performed (using a custom python script) to exclude any record whose genome sequence: (i) is sampled from a non-human host; (ii) has a length shorter than 29,000⍰nt; (iii) lacks a complete sample collection date (i-e YYYY-MM-DD); and (iv) was not collected between the start and end date provided in the command (by default no limits). Following that, the filtered metadata file and the sequence file are split into multiple files (1 for each lineage). Sequence files are then directed to BBMap [22] for further quality control. Sequences with more than 580 (∼2%) uncalled bases (Ns) are filtered in this process. The filtered sequences are then directed to the minimap2 (default; recommended) [23] or BWA-MEM [24] as two alternate routes for pairwise alignment against the SARS-CoV-2 reference genome (GenBank accession MN908947.3) [4]. The resulting BAM (compressed binary version of a sequence alignment file) is directed to call variants using one of Freebayes (default; recommended) [25] or Ivar [26] variant calling tools. Freebayes produces a Genomic Variant Call Format (GVCF) which is further processed using a custom Python script to filter low-quality mutations and produce a Variant Call Format file (VCF).

#### 2.2.3 Annotation

There is a multi-step annotation process for the mutations in the VCF file. The first step is to flag problematic sites using the curated list maintained at https://github.com/W-L/ProblematicSites_SARS-CoV2 (last accessed May 15th, 2022) [27]. The process is done using a custom Python script. The next step is to profile each mutation in the VCF file. SnpEff [28] is to profile (i) mutation type (e.g., Single Nucleotide Polymorphism (SNP), Insertion (INS), etc.); (ii) effect of the mutation in severity (e.g., High, Moderate, Low, Modifier); (iii) position of the mutation (e.g., gene name, upstream or downstream of a gene); and (iv) amino acid and codon change. The updated file is directed to the next phase where the mutations identified in the region of mature peptides are annotated with respective mature peptides. A custom Python script uses SARS-CoV-2 reference genome annotation available at NCBI [29] and extracts mature peptide features, and maps it to mutations based on the position.

##### 2.2.3.1 Pokay

We have developed a manually curated standardized resource to capture the functional annotation from the available literature for SARS-COV-2 mutations. The data is being captured in a hierarchical structure, where different functional categories are paired with genes e.g., *S_convalescent_plasma_escape* describes the function(s) related to convalescent plasma escape in the S gene (spike protein). Within these categories, specific functions are captured e.g., *Decreases in neutralization efficiency up to ∼25-50% in the S2D97 antibiotic*. Each function has a mutation or list of mutations that have been studied together to have a functional impact. Finally, for every function, additional information including the single or multiple source(s), citation(s), and URL(s) are listed for each study related to the function e.g., *Thomson et al. (2020)* https://www.biorxiv.org/content/10.1101/2020.11.04.355842v1. This structure of data organization allows us to disentangle complex relations between mutations and functions. A mutation can be involved in single or multiple functions, similarly, a functional impact can be associated with a single mutation, multiple individual mutations, or multiple mutations in a combined form. A custom python script is developed to merge the functional data from all functional categories to generate a combined file that is integrated into the genomics variant called data for visualization and surveillance reports (see Surveillance reports). Currently, in v.0.4.0 we have curated more than 70 unique function categories and over 4000 functional annotations for at least 1000 mutations either individually or in combination. This format also allows the user to provide SARS-CoV-2 functional annotations from a different source as long as the format is compatible. A webpage is autogenerated from the pokay repository for an interactive view of functions and mutations curated https://people.ucalgary.ca/~gordonp/front.html. Efforts are being made to regularly update pokay and release monthly versions with the latest information about mutations and their associated functions through extensive literature search and curation.

#### 2.2.4 Surveillance reports

The annotation process generates a GVF file that is used for visualization. The GVF file also undergoes a series of steps to be summarized into a surveillance report. Two types of surveillance reports are generated: (i) A TSV file that contains each identified mutation and its corresponding functions in a row along with contextual information; and (ii) a PDF file that contains summarized information in the form of tables. The TSV file can be used as an input for integrating COVID-MVP with other surveillance and downstream analysis tools. The description, type, and source of each column in the TSV file are available at the *nf-ncov-voc* GitHub repository (see the *Availability of Data and Material* section). The summarized PDF surveillance file includes three sections: the first section *Information on the variant or cluster of sequences section* includes (i) the range of dates for genomes based on the metadata field “*sample collection dates*” that are either provided by the user or extracted from the dataset; (ii) the number of genomes used in the analysis, to give context for the frequencies provided in section 3; and (iii) a list of Pangolin lineages identified in the dataset. The second section *(indicator section)* contains a table of three columns that provide the key indicators for public health surveillance, e.g. *transmissibility between humans, infection severity, etc*. (indicators can be specified for each run in an input file: see the *Availability of Data and Material* section), functional categories manually curated in functional annotation resource (as discussed above), categorized among the indicators (e.g. *ACE2 receptor binding affinity, viral load*, and *outcome hazard ratio* are classified under *infection severity)* and the mutations identified for each category and indicator. Finally, the third section (*mutation significance section)* consists of a multi-column table that provides the prevalence, functional impact of each mutation, hyperlinked citation for the study, lineage, number of sequences in the lineage from the dataset, reference allele, alternate allele, and alternate frequency. An example of a surveillance report in PDF format is available in the *nf-ncov-voc* repository (see the *Availability of Data and Material* section).

## 2.3 Deployment

We have deployed COVID-MVP on a cloud service and have developed a template for users to follow. We used a gunicorn server [30] to serve the python-based frond-end application from a docker container [31], with the image built with docker-compose. The docker image for the application is built on an anaconda base image for docker which provides essential packages for the functioning of the application. For establishing a secure SSL connection, we used an Nginx reverse proxy setup [32]. The combination of this framework provides the ability to run a multi-threading server through gunicorn. It also optimizes workers to handle varying levels of traffic. Containerization with docker increases the portability of the application by providing the required dependencies preinstalled. Finally, Nginx reverse proxy provides a means for easy and secure access to the application. Successful deployment on a Linux server will require user access with ***sudo*** permissions. Template repository is available as an open-source project which can be cloned to deploy COVID-MVP. See the *Availability of Data and Material* section.

## 3. Discussion

COVID-19 surveillance and reporting have started to slow down despite the uptick in a highly infectious subvariants of Omicron including BA.2. Case rates and hospitalizations are on the rise again [33]. Due to the extensive SARS-CoV-2 genomic sequencing, several tools, workflows, and/or visualization frameworks have been developed for surveillance and research purpose. Most of these tools have added value in different ways. We have developed a combination of visualization and analytical workflow to provide the functional impact of the mutations identified in SARS-CoV-2 variants of concern and variants of interest. COVID-MVP underscores the importance of tracking the functional impact of mutations in the already circulating and emerging lineages. These functions are manually curated hierarchically in the POKAY repository. Where literature is available, each mutation has a function associated with it and is classified under a functional category. We also classify these functional categories into public health surveillance indicators to create a summarized functional report. Like CovMT [17], COVID-MVP also provides the prevalence of mutations in the visualization and surveillance report. However, since COVID-MVP and nf-ncov-voc can be used offline as well, reports can be summarized from different datasets, which is especially useful in cases where data is not public yet (e.g., data from public health labs) (see Supp Table 1). Moreover, due to the modular and reproducible nature of the analytical workflow, the public health surveillance indicators can be updated based on the changing epidemiological parameters as we navigate through different phases of the pandemic. COVID-MVP can also be used for visualizing and analyzing data from environmental surveillance of SARS-CoV-2 such as wastewater surveillance. Even though our analytical workflow was initially conceived to use whole-genome sequencing data, we added different entrance points into the nextflow-based workflow to allow the processing of different types of data inputs. Environmental surveillance primarily is done using metagenomic sequencing [34–38]. As metagenomic sequencing is a non-targeted sequencing, it requires additional bioinformatics steps (not included in the workflow) to remove non-SARS-CoV-2 sequences before variant calling. Both the frontend visualization and analytical workflow can take a user-provided VCF as an input. In this case, as the workflow detects the file type of input data and runs a required set of steps, the first two sub-workflows are skipped and data is directed into the annotation sub-workflow (see fig. 3). This flexibility of workflow and visualization makes COVID-MVP adaptable for different SARS-CoV-2 surveillance strategies. The COVID-19 pandemic has also highlighted the bioinformatics bottleneck that exists specifically in analytical workflows and visualization tools [13]. It has been established those analytical workflows and visualizations must be developed in a structure rapidly adaptable to other pathogens. COVID-MVP’s standalone visualization application and modular genomic workflow make sure that it can be adapted to other circulating and emerging pathogens such as *Vibrio* or *Influenza* with structured updates.

## 4. Future Work

We aim to continue developing and adding features to COVID-MVP visualization based on the evolution of SAS-CoV-2. We also intend to expand this visualization framework to other circulating pathogens. Similarly, we are curating functional annotations as the literature grows for SARS-CoV-2 mutations. Additionally, to ensure maximum reusability, we are working to ontologize the functional categories and their descriptions from Pokay. This will result in a standardized framework for describing the functional impact of mutations, which can be reused in the future for genomic surveillance of other pathogens.

## 5. Conclusion/Concluding Remarks

Since the start of the SARS-CoV-2 pandemic, it has been established that molecular surveillance played a crucial role in tracking the evolution and spread of SARS-CoV-2. This has been made possible due to the remarkable amount of genomic sequencing and its availability through global or local databases such as GISAID, the COVID-19 Data Portal at NCBI, European Bioinformatics Institute (EBI), or Canadian VirusSeq Data Portal. Several web-based and offline tools have been developed to process these datasets and provide information primarily on the SARS-CoV-2 lineages and variants using phylogenetic reconstruction. We believe that COVID-MVP provides additional value in connecting the mutations with their biological impact. It flags the mutations that have been identified in various lineages ad sub-lineages that may have a significant effect on the biology of the virus. COVID-MVP is a hybrid application that can be used as a web-based dashboard or offline surveillance tool in combination with the genomics workflow (nf-ncov-voc), to enhance molecular surveillance at a mutation level. COVID-MVP will support SARS-CoV-2 surveillance and basic research by providing mutation profiles and functional impacts through interactive visualization and summarized reports.

## Abbreviations

COVID-MVP: Coronavirus - Mutations, Variants, and Prevalence
DEL: Deletion
GISAID: Global Initiative on Sharing Avian Influenza Data
GVCF: Genomic Variant Call Format
GVF: Genome Variant File
INS: Insertion
SNP: Single Nucleotide Polymorphism
VCF: Variant Called File
VOC: Variants of Concern
VOI: Variants of Interest
VUM: Variants Under Monitoring
WGS: Whole Genome Sequencing
WHO: World Health Organization

## Declarations

### Ethics approval and consent to participate

Not Applicable

### Availability of data and materials

1. COVID-MVP running instance - https://covidmvp.cidgoh.ca
2. COVID-MVP source code and manual - https://github.com/cidgoh/COVID-MVP
3. COVID-MVP deployment source code - https://github.com/cidgoh/covidmvp_deployment
4. nf-ncov-voc source code - https://github.com/cidgoh/nf-ncov-voc
5. Pokay - https://github.com/nodrogluap/pokay
6. SARS-CoV-2 surveillance indicators for public health - https://github.com/cidgoh/nf-ncov-voc/tree/master/assets/ncov_surveillanceIndicators
7. Surveillance report (PDF format example) - https://github.com/cidgoh/nf-ncov-voc/blob/master/docs/Omicron_surveillance_report.pdf
8. All sequence data used in developing, testing, and generating the figure is available at https://virusseq-dataportal.ca

## Authors’ contributions

All authors reviewed the manuscript. M.Z.A led and developed the backend Nextflow workflow with regular feedback from all authors. M.Z.A co-implemented the Python + LaTeX code to summarize mutation data and generate surveillance reports. M.Z.A co-drafted the manuscript. I.S.G led and developed the front-end component with regular feedback from all the authors. I.S.G co-drafted the manuscript. M.I implemented the Python code to format mutation data and integrated it with functional annotation. M.I co-implemented the Python + LaTeX code to summarize mutation data and generate surveillance reports. M.I contributed to the curation of functional annotations in Pokay. A.S implemented the Python code to process functional annotation data curated in Pokay and contributed to Pokay data curation. K.V contributed to the curation of functional annotations in Pokay. K.D.I deployed the COVID-MVP public instance and developed a template for users to do at their shared or individual systems based on system permissions. P.G developed the Pokay tool and curated functional annotations. P.G provided regular feedback on genomic data analyses and visualization components. G.V.D provided the template for the surveillance report in addition to regular feedback on the visualization, genomics data analysis, and the manuscript. W.W.H conceived, administered, and supervised the project. W.W.H reviewed and edited the manuscript. W.W.H acquired the funding.

## Acknowledgments

The results here are in whole, or part based upon data hosted at the Canadian VirusSeq Data Portal: https://virusseq-dataportal.ca/. We wish to acknowledge the Canadian Public Health Laboratory Network (CPHLN), Genome Canada, and the CanCOGeN VirusSeq Consortium for their contribution to the Portal, see supplementary file for detailed information

