## Supplementary File for "COVID-MVP: an interactive visualization for tracking SARS-CoV-2 mutations, variants, and prevalence, enabled by curated functional annotations and portable genomics workflow"

Supplementary File 01:

### Supplementary Information 1

|  | Tracking Mutations | Functional Annotation | Ability to upload sequences | Summarized surveillance reports | Ability to view data geographically | Longitudinal analysis | Ability to download and use as local application |
| --- | --- | --- | --- | --- | --- | --- | --- |
| CovMT | ✓ | ❌ | ✓ | ❌ | ✓ | ✓ | ❌ |
| CovRadar | S-Gene only | ❌ | ❌ | ❌ | ✓ | ✓ | ❌ |
| Outbreak.info | ✓ | ❌ | ❌ | ✓ | ✓ | ✓ | ❌ |
| **COVID-MVP** | **✓** | **✓** | **✓** | **✓** | **✓** | **✓** | **✓** |

Supplementary Table 1. Comparison of COVID-MVP with similar SARS-CoV-2 mutation tracking dashboards/tools

### **Acknowledgements**

Canadian Public Health Laboratory Network (CPHLN) members and staff have contributed data to the portal

**Alberta**

*Genomes with prefix ABPHL*

Buss, E, Croxen M, Deo A, Dieu P, Gill K, Ferrato C, Khan F, Koleva P, Li V, Lloyd C, Lynch T, Ma R, Murphy S, Pabbaraju K, Shokoples S, Tipples G, Thayer J, Whitehouse M, Wong A, Yu C, Zelyas N

*Genomes in collaboration with UC (prefix AB-NNNNNN)*

Gordon P, Lam LG, Pabbaraju K, Wong A, Ma R, Li V, Melin A, Tipples G, Berenger B, Zelyas N, Kellner J, Bernier F, Chui L, Croxen M

**British Columbia**

*BCCDC Public Health Laboratory:*

Prystajecky Natalie, Linda Hoang, John R. Tyson, Dan Fornika, Shannon Russell, Kim MacDonald, Kimia Kamelian, Ana Pacagnella, Corrinne Ng, Loretta Janz, Richard Harrigan, Robert Azana, Mel Krajden

**Manitoba**

*Cadham Provincial Laboratory collected specimens sequenced at the National Microbiology Lab (NML):*

Paul Van Caeseele, Jared Bullard, David Alexander, Kerry Dust.

*NML:* Anna Majer, Shari Tyson, Grace Seo, Philip Mabon, Elsie Grudeski, Rhiannon Huzarewich, Russell Mandes, Anneliese Landgraff, Jennifer Tanner, Natalie Knox, Morag Graham, Gary Van Domselaar, Nathalie Bastien, Yan Li, Timothy Booth, Darian Hole, Madison Chapel, Kirsten Biggar

*Cadham Provincial Laboratory sequenced specimens:*

David Alexander, Lori Johnson, Janna Holowick, Joanne Sanders, Adam Hedley, Kerry Dust

*Dynacare sequenced specimens:*

Hilary Racher, Melissa Desaulnier, Tintu Abraham, Hongbin Li (Impact Genetics, Brampton Ontario)

**New Brunswick**

Centre Hospitalier Universitaire Georges L. Dumont; National Microbiology Laboratory

Richard Garceau, Guillaume Desnoyers, Nicolas Crapoulet, Pierre Lyons, Woodson Shaw and Simi Chacko, Anna Majer, Shari Tyson, Grace Seo, Philip Mabon, Elsie Grudeski, Rhiannon Huzarewich, Russell Mandes, Anneliese Landgraff, Jennifer Tanner, Natalie Knox, Morag Graham, Gary Van Domselaar, Nathalie Bastien, Yan Li, Timothy Booth, Darian Hole, Madison Chapel, Kirsten Biggar.

**Newfoundland and Labrador**

Dr. Leonard A. Miller Centre for Health Services; National Microbiology Laboratory

Robert Needle, Yang Yu, Laura Gilbert, George Zahariadis, Chris Corkum, Anna Majer, Shari Tyson, Grace Seo, Philip Mabon, Elsie Grudeski, Rhiannon Huzarewich, Russell Mandes, Anneliese Landgraff, Jennifer Tanner, Natalie Knox, Morag Graham, Gary Van Domselaar, Nathalie Bastien, Yan Li, Timothy Booth, Darian Hole, Madison Chapel, Kirsten Biggar, Kerri Smith, Matthew Gilmour.

**Nova Scotia**

*QEII Health Sciences Centre*; National Microbiology Laboratory*

Todd Hatchette, Jason LeBlanc, Janice Pettipas, Dan Gaston, Greg McCracken, Nathalie Bastien, Yan Li, Timothy Booth, Darian Hole, Madison Chapel, Kirsten Biggar, Anna Majer, Shari Tyson, Grace Seo, Philip Mabon, Elsie Grudeski, Rhiannon Huzarewich, Russell Mandes, Anneliese Landgraff, Jennifer Tanner, Natalie Knox, Morag Graham, Gary Van Domselaar.

(*data tagged post-September 14th should include Allana Loder)

**Ontario**

*Public Health Ontario Laboratory*

Vanessa G Allen, Philip Banh, Yao Chen, Alireza Eshaghi, Nahuel Fittipaldi, Christine Frantz, Jonathan B Gubbay, Jennifer L Guthrie, Esha Joshi, Aimin Li, Michael C.Y. Li, Dean Maxwell, Sandeep Nagra, Samir N. Patel, Karthikeyan Sivaraman, Ashleigh Sullivan, Sarah Teatero, Andre Villegas, Matthew Watson, Sandra Zittermann

*Ontario Institute for Cancer Research*

Jared T. Simpson, Richard de Borja, Paul Krzyzanowski, Bernard Lam, Lawrence Heisler, Michael Laszloffy, Yogi Sundaravadanam, Ilinca Lungu, Lubaina Kothari, Cassandra Bergwerff, Jeremy Johns, Felicia Vincelli, Philip Zuzarte

*McMaster University*

Hooman Derakhshani, Sheridan J.C. Baker, Emily M. Panousis, Ahmed N. Draia, Jalees A. Nasir, Michael G. Surette, Andrew G. McArthur

**Québec**

*Laboratoire de Santé Publique du Québec, McGill Génome Sciences Centre; CoVSeq Consortium*

Sandrine Moreira, Jiannis Ragoussis, Guillaume Bourque, Jesse Shapiro, Éric Fournier, Réjean Dion, Hugues Charest, Aurélie Guilbault, Benjamin Delisle, Sarah Reiling, Anne-Marie Roy, Shu-Huang Chen, Corinne Darmond, Sally Lee, Brent Brookes, Pierre Lepage, Jannick St-Cyr, Patrick Willet, Mathieu Bourgey, David Bujold, Hector Galvez, Paul Stretenowich, Pierre-Olivier Quirion, Romain Grégoire, Carmen Lia Murall, Julie Hussin, Raphaël Poujol, Jean-Christophe Grenier, Fatima Mostefai, Sylvie Laboissières, Alexandre Montpetit, Mark Lathrop, Michel Roger

**Saskatchewan**

*Roy Romanow Provincial Laboratory*

Ryan McDonald, Keith MacKenzie, Meredith Faires, Kara Loos, Stefani Kary, Rachel DePaulo, Laura Klassen, Alanna Senecal, Amanda Lang, Jessica Minion, Roy Romanow Provincial Laboratory - Molecular Diagnostics.

*National Microbiology Laboratory*

Anna Majer, Shari Tyson, Grace Seo, Philip Mabon, Elsie Grudeski, Rhiannon Huzarewich, Russell Mandes, Anneliese Landgraff, Jennifer Tanner, Natalie Knox, Morag Graham, Gary Van Domselaar, Nathalie Bastien, Yan Li, Timothy Booth, Darian Hole, Madison Chapel, Kristen Biggar, Emily Haidl, Chanchal Yadav, Jeff Tuff, Connor Chato, Katherine Eaton, and Adrian Zetner.

**Academic, Health Network and Research Institutions staff have contributed data to the portal**

*Unity Health Toronto*

Ramzi Fattouh, Larissa M. Matukas, Yan Chen, Mark Downing, Trina Otterman, Karel Boissinot, Le Luu

*University Health Network/Mount Sinai Hospital Department of Microbiology*

Marie-Ming Aynaud, Javier Hernandez, Seda Barutcu, Kin Chan, Jessica Bourke, Marc Mazzulli, Tony Mazzulli, Laurence Pelletier, Jeff Wrana, Aimee Paterson, Angel Liu, Allison McGeer

*Kingston Health Sciences Centre and Queen’s University*

Prameet M. Sheth, Calvin Sjaarda, Robert Colautti, Katya Douchant

*University of British Columbia*

John R. Tyson, Gabrielle Jayme, Karen Jones, Terrance P. Snutch

*Toronto Invasive Bacterial Diseases Network; Sunnybrook Health Sciences*

Allison McGeer, Patryk Aftanas, Angel Li, Kuganya Nirmalarajah, Emily Panousis, Ahmed Draia, Jalees Nasir, David Richardson, Michael Surette, Samira Mubareka, Andrew G. McArthur

*Eastern Ontario Regional Laboratory Association*

Leanne Mortimer, Hooman Derakhshani, Emily Panousis, Ahmed Draia, Jalees Nasir, Robert Slinger, Andrew G. McArthur

**CanCOGeN VirusSeq committees and working groups’ members**

*CanCOGeN VirusSeq Implementation Committee*

Terrance Snutch, Fiona Brinkman, Marceline Côté, William Hsiao, Yann Joly, Sharmistha Mishra, Sandrine Moreira, Samira Mubareka, Jared Simpson, Megan Smallwood, Gary Van Domselaar

*CanCOGeN Capacity Building Working Group*

Gary Van Domselaar, Matthew Croxen, Natalie Knox, Celine Nadon, Jennifer Tanner

*CanCOGeN Data Analytics Working Group*

Gary Van Domselaar, Fiona Brinkman, Zohaib Anwar, Robert Beiko, Matieu Bourgey, Guillaume Bourque, Richard de Borja, Ahmed Draia, Jun Duan, Marc Fiume, Dan Fornika, Eric Fournier, Erin Gill, Paul Gordon, Emma Griffiths, Jose Hector Galvez Lopez, Darian Hole, William Hsiao, Jeffrey Joy, Kimia Kamelian, Natalie Knox, Philip Mabon, Finlay Maguire, Tom Matthews, Andrew McArthur, Samir Mechai, Sandrine Moreira, Art Poon, Amos Raphenya, Claire Sevenhuysen, Jared Simpson, Jennifer Tanner, Lauren Tindale, John Tyson, Geoff Winsor, Nolan Woods

*CanCOGeN Ethics and Governance Working Group*

Yann Joly, Fiona Brinkman, Erin Gill, William Hsiao, Hanshi Liu, Sandrine Moreira, Gary Van Domselaar, Ma’n Zawati, Sarah Savić-Kallesøe

*CanCOGeN Metadata Working Group*

William Hsiao, David Alexander, Zohaib Anwar, Nathalie Bastien, Tim Booth, Guillaume Bourque, Fiona Brinkman, Hughes Charest, Caroline Colijn, Matthew Croxen, Guillaume Desnoyers, Rejean Dion, Damion Dooley, Ana Duggan, Leah Dupasquier, Kerry Dust, Eleni Galanis, Emma Garlock, Erin Gill, Gurinder Gopal, Tom Graefenhan, Morag Graham, Emma Griffiths, Linda Hoang, Naveed Janjua, Jeffrey Joy, Kimia Kamelian, Lev Kearney, Natalie Knox, Theodore Kuschak, Jason LeBlanc, Yan Li, Anna Majer, Adel Malek, Ryan McDonald, David Moore, Celine Nadon, Samir Patel, Natalie Prystajecky, Anoosha Sehar, Claire Sevenhuysen, Garrett Sorensen, Laura Steven, Lori Strudwick, Marsha Taylor, Shane Thiessen, Gary Van Domselaar, Adrian Zetner

*CanCOGeN Research Collaborations Working Group*

Fiona Brinkman, Zohaib Anwar, Marceline Côté, Marc Fiume, Laura Gilbert, Erin Gill, Paul Gordon, Yann Joly, Sandrine Moreira, Samira Mubareka, Natalie Prystajecky, Jennifer Tanner, Gary Van Domselaar, Phot Zahariadis,

*CanCOGeN Sequencing Working Group*

Ioannis Ragoussis, Terrance Snutch, Patryk Aftanas, Matthew Croxen, Hooman Derakhshani, Nahuel Fittipaldi, Morag Graham, Andrew McArthur, Sandrine Moreira, Samira Mubareka, Natalie Prystajecky, Ioannis Ragoussis, Jared Simpson, Michael Surette, John Tyson

*CanCOGeN Quality Control Working Group*

Jared Simpson, Mathieu Bourgey, Kodjovi Dodji Mlaga, Nahuel Fittipaldi, Jose Hector Galvez Lopez, Natalie Knox, Genevieve Labbe, Pierre Lyons, Philip Mabon, Finlay Maguire, Anna Majer, Andrew McArthur, Ryan McDonald, Sandrine Moreira, Natalie Prystajecky, Karthikeyan Sivaraman, Kerri Smith, Terrance Snutch, Karthikeyan Sivaraman Andrea Tyler, John Tyson, Gary Van Domselaar

*Canadian VirusSeq Data Portal (CVDP) Team*

Guillaume Bourque, Lincoln Stein, Christina Yung, Hanshi Liu, Yann Joly, Adrielle Houweling, William Hsiao, Marc Fiume, David Bujold, Erin Gill, Fiona Brinkman, Nithu John, Rosita Bajari, Linda Xiang, Alexandru Lepsa, Jaser Uddin, Justin Richardson

Funding for the VirusSeq Data Portal is provided by The Canadian COVID Genomics Network (CanCOGeN), and supported by Genome Canada and Innovation, Science and Economic Development Canada (ISED)
